# Real-time visualization of spatial muscle activity using augmented reality improves endurance during dynamic knee extensions

**DOI:** 10.64898/2026.11.11.765967

**Authors:** Michail Arvanitidis, Giacinto Luigi Cerone, Marco Gazzoni, Deborah Falla

**Affiliations:** Centre of Precision Rehabilitation for Spinal Pain (CPR Spine), School of Sport, Exercise and Rehabilitation Sciences, College of Life and Environmental Sciences, University of Birmingham, Birmingham, UK; Laboratory for Engineering of the Neuromuscular System (LISiN) and PoliToBIOMed Laboratory, Department of Electronics and Telecommunications, Politecnico di Torino, Turin, Italy

**Keywords:** Augmented reality, Biofeedback, High-density electromyography, Muscle fatigue, Spatial distribution

## Abstract

**Purpose:** High-density surface electromyography (HDsEMG) quantifies the spatial distribution of muscle activity, and greater redistribution during sustained muscle contractions has been associated with longer endurance time. However, it remains unclear whether providing real-time biofeedback on the spatial distribution of muscle activity can enhance endurance. This study determined whether real-time augmented reality (AR) biofeedback on the spatial distribution of knee extensor muscle activity allowed individuals to alter the spatial activation of their muscles during dynamic contractions and improved knee extensor endurance.

**Methods:** Fourteen healthy adults performed isokinetic knee extensions at 35% of maximal voluntary contraction torque until volitional exhaustion under two randomized conditions: with and without AR HDsEMG biofeedback. HDsEMG was recorded from the vastus lateralis and rectus femoris using a 64-electrode grid.

**Results:** Participants completed more repetitions with biofeedback than without (*p < 0.001*, *d = 1.41*). With biofeedback, a lateral shift in the spatial distribution of muscle activity was observed (medio-lateral centroid; *p = 0.022*), indicating greater contribution from the lateral region of the quadriceps. Global markers of myoelectric manifestations of fatigue showed similar temporal changes in both conditions. Exploratory analyses showed no association between condition-related changes in HDsEMG variables and the individual endurance gain (all *p > 0.05*).

**Conclusions:** AR biofeedback based on HDsEMG can improve muscular endurance while enabling individuals to modify spatial activation of their knee extensors during dynamic fatiguing exercise. These findings support the use of AR HDsEMG biofeedback as a novel tool to optimize neuromuscular function and potentially push the boundaries of athletic performance.

## 1. INTRODUCTION

High-density surface electromyography (HDsEMG) is an electrophysiological technique that can be used to quantify the spatial distribution of activity within muscles across a large portion of the muscle surface. While intramuscular EMG and conventional bipolar surface electromyography (EMG) provide spatially localized information on muscle activation, HDsEMG uses grids of closely spaced electrodes covering a large portion of the muscle surface to capture the spatial summation of motor unit (MU) action potentials, enabling a topographical representation of muscle activation. This allows the assessment of regional differences in activation, typically described using the centroid (barycenter) of the EMG amplitude distribution and measures that quantify the homogeneity of the activation distribution across the grid, such as modified entropy.

Studies using HDsEMG have consistently shown that muscle activity is not uniformly distributed within muscles (1–4). In healthy individuals, sustained or repetitive contractions are accompanied by a progressive redistribution of activity across muscle regions, likely reflecting a strategy to delay local fatigue by progressively recruiting MUs in different regions while reducing the activity of those engaged since the onset of the contraction (5). In contrast, redistribution of muscle activity is typically diminished in individuals with musculoskeletal pain (6–9). For example, people with musculoskeletal conditions such as chronic low back pain or fibromyalgia commonly exhibit more fixed activation patterns, with activity concentrated in a smaller region of the muscle (i.e., a more focal distribution), during both isometric and dynamic tasks (5, 6, 9–12). This may be clinically meaningful, since the degree of spatial redistribution is known to relate to endurance performance; greater shifts in activation are strongly associated with a longer time to task failure (13, 14). Therefore, this more focal activation and reduced capacity to redistribute load between muscle regions may contribute directly to localized overload, earlier fatigue, and symptom persistence (5).

Collectively, these findings provide a rationale for using HDsEMG biofeedback to promote variability in muscle activation. In an earlier study, HDsEMG biofeedback of muscle activity was displayed on an external computer monitor during an isometric task targeting the upper trapezius; although participants could modify their activation pattern with the use of HDsEMG biofeedback, this did not translate into improved endurance (15). Several factors likely constrained the performance outcomes in that study. Primarily, the isometric nature of the task and the relatively limited volume of the upper trapezius assessed may have restricted meaningful load redistribution between regions of the muscle. Addressing these constraints requires investigating biofeedback during a dynamic task involving a larger, more architecturally complex muscle group, while also overcoming a practical limitation of screen-based feedback systems, namely that participants must repeatedly look away from the task toward an external monitor and mentally map an abstract two-dimensional image onto their own body.

The vastus lateralis (VL) exhibits regional heterogeneity in its internal architecture, with differences in fascicle length and pennation angle between the middle and distal portions that influence regional mechanical behavior during dynamic isokinetic contractions (16). At the neural level, the activity of VL MUs is represented by multiple independent latent factors (17). Moreover, the MU pools of the vasti share common synaptic inputs, with some MUs in one muscle being driven by the dominant neural module of the synergist (18), providing a neuroanatomical basis for the coordinated redistribution of activity across quadriceps compartments.

Whether these architectural and neurally distinct regions are volitionally accessible remains an open question in motor control research. Recent MU decomposition studies suggest that voluntary dissociation of activity between quadriceps compartments is achievable when real-time intramuscular EMG feedback is provided during low-force, non-fatiguing isometric contractions (19). However, whether voluntary alteration of the spatial distribution of muscle activity, as captured by HDsEMG, can be achieved during dynamic fatiguing tasks, and whether such modulation influences endurance performance, has not been investigated. To solve the visual constraints of previous setups, we have developed an augmented reality (AR) application that integrates directly with a wireless HDsEMG acquisition system (20). This system allows participants to continuously monitor the spatial distribution of their muscle activity since it projects real-time topographical maps of the EMG amplitude directly over the electrode grid.

Therefore, the aim of this study was to examine the effects of real-time AR HDsEMG biofeedback during dynamic, fatiguing knee extensions in healthy individuals. We hypothesized that individuals will be able to use the AR HDsEMG biofeedback in a meaningful way to alter their muscle activation pattern, and as a result, change the spatial distribution of muscle activity resulting in improved endurance.

## 2. METHODS

### 2.1. Study design and overview

A repeated-measures crossover design was implemented. Participants attended two laboratory sessions (with and without AR biofeedback) at the Centre of Precision Rehabilitation for Spinal Pain, University of Birmingham, United Kingdom, separated by a minimum washout period of one week to mitigate residual fatigue and learning effects. Condition (session) order was randomized and counterbalanced across participants.

All participants provided written informed consent prior to data collection. The study protocol was approved by the University of Birmingham Science, Technology, Engineering and Mathematics Ethical Review Committee (ERN_2022-0219) and was conducted in accordance with the Declaration of Helsinki. EMG reporting follows the CEDE-Check guidelines (21).

### 2.2. Participants

Fourteen healthy, recreationally active volunteers (aged 18–35 years) were recruited from the local community. The upper age limit was established to minimize potential confounding effects of age-related musculoskeletal changes on neuromuscular performance. Participants were excluded if they had any history of lower limb musculoskeletal or neurological conditions, including previous knee injury or surgery.

### 2.3. General Procedures

Demographic details were recorded and participants completed the International Physical Activity Questionnaire – Long Form (IPAQ-LF), a self-report instrument with established reliability and validity for assessing habitual physical activity levels (22). Subsequently, an isokinetic dynamometer was set up and individually adjusted for each participant (see Dynamometer setup), followed by identification and marking of electrode placement sites, skin preparation, and application of the HDsEMG electrodes (see Skin preparation and electrode placement). Participants were then given time to warm up and practice the knee extension task at the required speed and torque. In sessions involving the AR biofeedback condition, the biofeedback system and display were explained, and participants were given approximately 10 minutes to practice with the real-time HDsEMG topographical maps during low-level contractions, allowing them to learn how to modulate the spatial distribution of quadriceps muscle activation. The experimental protocol consisted of three dynamic maximum voluntary contractions (MVCs) of knee extension separated by 2 minutes of rest, followed by 10 minutes of rest, after which participants performed repeated submaximal contractions at 35% MVC until task failure. After every three repetitions, participants rated their perceived exertion on the Borg CR10 scale (23). Immediately upon task failure, participants performed one additional MVC to quantify the extent of exercise-induced fatigue.

### 2.4. Dynamometer setup

Participants were seated on an isokinetic dynamometer (Biodex System 3, Biodex Medical Systems, Shirley, NY) in an adjustable chair, with the trunk kept vertical and the hip, knee, and ankle joint angles maintained at 90°. The dynamometer’s rotational axis was aligned with the right lateral femoral epicondyle, and the lower leg was secured to the lever arm just above the lateral malleolus. Straps were used to stabilize the pelvis and upper torso to isolate the knee joint. All dynamometer settings were recorded for each participant to ensure identical positioning across sessions.

Then, participants performed three dynamic MVCs of concentric knee extension through the full range of motion (90° to 30° of knee flexion) at 10°/s. Each contraction lasted approximately 6 seconds, with 2 minutes of rest between attempts. The highest MVC value from the first session defined the submaximal target torque for both conditions to ensure the workload was exactly the same between the two sessions. The MVC from the second session was recorded to confirm there were no baseline differences in maximal strength across visits.

### 2.5. Skin preparation and electrode placement

Prior to electrode placement, the skin over the right thigh of all participants was shaved (if necessary), mildly abraded using an abrasive gel (Nuprep Skin Prep Gel, Weaver and Company, Aurora, CO), and cleansed with water to minimize impedance (24). Two adjacent semi-disposable 32-electrode grids (11 × 3 configuration each; 15-mm inter-electrode distance), forming a combined 11 × 6 (rows × columns) array of 64 electrodes (ReC Bioengineering and LISiN, Turin, Italy) were used to detect HDsEMG signals. Electrode cavities were filled with conductive gel (AC-CREAM; SPES Medica, Genoa, Italy), and a reference electrode (WhiteSensor WS; Ambu A/S, Ballerup, Denmark) was placed over the patella.

The grid was then placed over the VL with the electrode columns aligned parallel to the muscle fibers. To achieve this, the array was oriented at 20° along a reference line drawn from the anterior superior iliac spine to the lateral border of the patella, as previously described (25–27). Based on initial pilot testing, the fourth electrode row, counting from the proximal edge of the grid, was positioned at 50% of this reference line; this placement allowed for recording from a large portion of the VL in addition to the lateral region of the rectus femoris (RF). The position of the grid was marked on the skin using a permanent ink marker, and all distances were recorded to ensure consistent grid placement across the two experimental sessions.

### 2.6. Data acquisition

Monopolar HDsEMG signals were recorded using a wireless acquisition system (MEACS, ReC Bioengineering Laboratories S.r.l. and LISiN, Politecnico di Torino, Turin, Italy) (28). These signals were conditioned with a 10–500 Hz bandwidth and a gain of 192 ± 1 V/V, then digitized at a sampling frequency of 2048 Hz per channel with 16-bit resolution. Simultaneously, the analog torque output from the dynamometer was transmitted via a BNC cable to a custom force amplifier (DUEBio, OT Bioelettronica, Turin, Italy). This signal was transmitted via Bluetooth to the acquisition laptop, where both HDsEMG and torque data were synchronized and visualized in real time using the Bp software platform (ReC Bioengineering Laboratories S.r.l., Turin, Italy).

### 2.7. Torque feedback

A custom visual and auditory feedback system was integrated into the Bp software to enable participants to maintain the target torque. To facilitate an unobstructed view of the AR interface (described below) while allowing for rapid torque adjustments, the monitor showing the feedback bar was positioned on the floor at a distance of 1 m from the participant (Figure 1). This position was kept identical across both experimental sessions to ensure consistency. Participants were instructed to maintain their torque output at the seventh bar (35% MVC), which remained green when the target was met. Deviations of ±5% MVC, that was one bar above or below the target (i.e., 30% or 40% MVC), were indicated by the bar turning yellow, accompanied by a low-frequency auditory tone. In contrast, deviations exceeding ±5% MVC triggered a high-frequency auditory cue. This dual-modality feedback encouraged precise torque maintenance while allowing participants to shift their primary visual focus and attention to the AR biofeedback during the experimental session.

**Figure 1.**
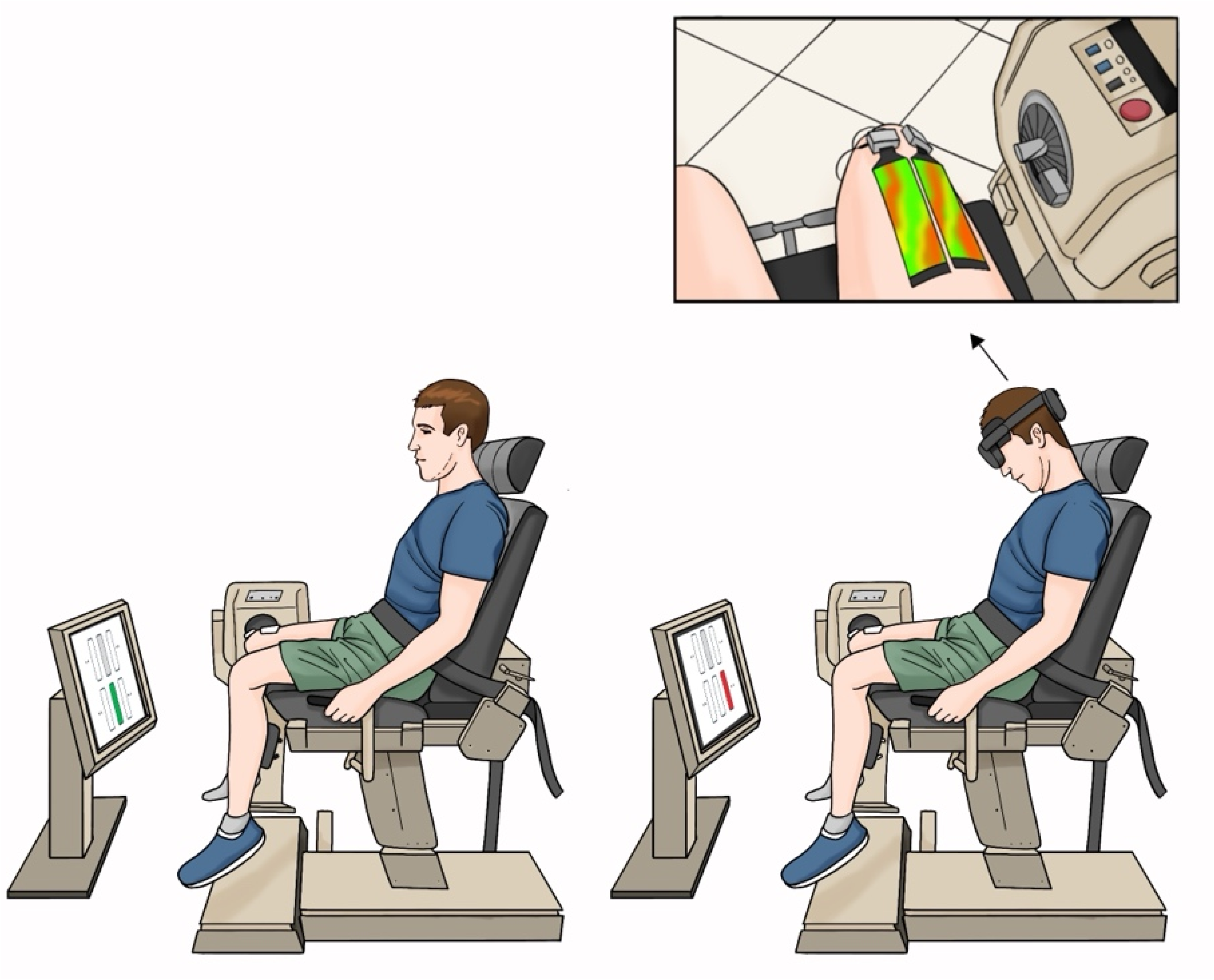
Experimental setup in the two conditions. Schematic of a participant seated on the isokinetic dynamometer performing dynamic knee extensions at 35% MVC without AR biofeedback (control condition; left) and with AR biofeedback (right). In both conditions, torque feedback was displayed on a monitor placed on the floor 1 m in front of the participant: the target bar (35% MVC) was green when torque output matched the target (left) and turned red when torque deviated by one bar (±5% MVC; shown here 5% above the target, right), accompanied by an auditory tone, so that participants could correct their torque output and return to the target. In the AR biofeedback condition, participants additionally wore a Microsoft HoloLens 2 head-mounted display, which projected a real-time topographical map of the RMS amplitude distribution recorded by the HDsEMG grid directly over the electrodes on the thigh (inset: the participant’s view through the headset). Warmer colors (red) indicate regions of higher and cooler colors (green, blue) regions of lower EMG amplitude.

### 2.8. Augmented reality biofeedback

Real-time AR biofeedback was provided via a Microsoft HoloLens 2 head-mounted display (Microsoft Corporation, Redmond, WA, USA), using a custom-developed application (MyoArt; ReC Bioengineering and LISiN, Politecnico di Torino, Turin, Italy). The interface presented a two-dimensional topographical map of the RMS amplitude distribution across the 11 × 6 electrode grid. This spatial map was color-coded to represent EMG magnitude, with warmer and cooler colors indicating higher and lower amplitudes, respectively. The AR overlay was projected directly into the participant’s field of view, allowing them to monitor the spatial distribution of their muscle activity without diverting their gaze from the torque feedback (Figure 1).

The same application also ran on a tablet, whose camera view of the participant’s thigh was overlaid with the same real-time topographical map, allowing the investigator to monitor the feedback from their own point of view. This was utilized to guide participants during familiarization and to ensure consistent engagement throughout the trial. During the biofeedback condition, participants were instructed to observe the topographical map and explore various activation strategies to modulate the spatial distribution of their muscle activity. No specific shift direction was suggested; instead, participants were encouraged to explore the spatial characteristics of their activation patterns. During the control condition, the task was performed without the AR headset, but participants were still provided with the torque biofeedback.

### 2.9. Signal processing

Signal processing was performed offline using custom MATLAB scripts (MathWorks, Natick, MA, USA). The monopolar signals were reorganized into their spatial configuration to form a 11×6 monopolar matrix. They were then visually inspected for noise and artefacts, and channels with poor signal quality were excluded (fewer than 5% of channels per recording). Single-differential signals were then computed longitudinally (row by row) within each grid to form a 10 × 6 differential matrix (58 channels, as the two positions without electrodes yielded no differential signal), which attenuated common-mode noise. These differential signals were bandpass filtered between 20 and 400 Hz using a fourth-order, zero-phase Butterworth filter.

For each repetition, the steady-state portion of the concentric phase was manually identified from the torque trace. Repetitions where the participant failed to sustain the target torque above 30% MVC for at least 2 s were excluded. For each channel and valid repetition, the root mean square (RMS) amplitude and mean frequency (MNF) were computed within non-overlapping 250-ms windows. MNF was estimated using the periodogram method with a rectangular window. These windowed values were then averaged to yield a single representative RMS and MNF value per channel for each repetition.

Obtaining one mean RMS value for each valid differential channel (up to 58) allowed for the generation of a 10 × 6 spatial EMG amplitude map per repetition. The spatial distribution of muscle activity was characterized from these EMG amplitude maps by calculating the centroid (barycenter) and modified entropy. The x-axis (medial–lateral) and y-axis (proximal–distal) centroid coordinates were computed as the weighted average of the electrode coordinates, using the RMS amplitude at each electrode as the weighting factor. Coordinates were expressed in millimeters relative to the most proximal electrode of the lateral-most column of the grid, with x increasing in the medial and y in the distal direction. Furthermore, modified entropy was calculated from the squared RMS values to quantify the heterogeneity of the spatial activation; higher entropy values indicate a more uniform distribution, whereas lower values reflect a more focalized pattern of muscle activation.

The torque signal was low-pass filtered at 15 Hz (third-order, zero-phase Butterworth filter) and demeaned using the first 2048 samples as a baseline reference. Raw torque values were converted from volts to N·m using a calibration factor derived from the dynamometer’s known output. Peak torque was identified from the pre-task and post-task MVCs. For each submaximal repetition, mean torque and the coefficient of variation (CoV = [standard deviation / mean] × 100) were calculated to assess torque steadiness and verify that mechanical output remained comparable across conditions.

Task performance was quantified using several parameters extracted from the torque signal: the total number of repetitions completed prior to task failure, the estimated total contraction time (number of repetitions × 6 s), and the total active time (sum of the durations of the individual contractions, identified manually from the torque trace as described above). Total session time was defined as the elapsed time from the onset of the first repetition to the completion of the last repetition. This allowed for the calculation of the average rest time per repetition [(total session time – total active time) / number of repetitions]. Although these performance measures are inter-related, all were computed for completeness and, in particular, to verify that pacing (the rest taken between repetitions) did not differ between conditions. Finally, to track the progression of fatigue over the same absolute workload in the two conditions, both sessions of each participant were truncated to the number of repetitions completed in the shorter session (in all cases, the session without biofeedback). These repetitions were divided into five consecutive epochs (E1–E5) containing an equal number of repetitions, and the HDsEMG and torque variables were averaged across the repetitions within each epoch. Consequently, E5 corresponded to task failure in the control condition but to a submaximal stage of the task in the biofeedback condition, so that between-condition differences reflect the response to an identical workload rather than differences in endurance time.

### 2.10. Statistical analysis

Statistical analyses were performed using SPSS (IBM SPSS Statistics, Version 30 for macOS, Armonk, NY). The exploratory correlation analyses, including the Holm adjustment and the correlation matrix (Supplementary Material), were performed in R (version 4.6.0; R Foundation for Statistical Computing, Vienna, Austria) using the *cor.test* and *p.adjust* functions of the stats package. Data distribution was first assessed using the Shapiro–Wilk test together with visual inspection of Q–Q plots to confirm that the assumptions for parametric testing were met. Descriptive data are presented as mean ± SD, except for self-reported physical activity (IPAQ-LF), which is presented as median (interquartile range, IQR) owing to its skewed distribution.

Performance outcomes (total number of repetitions, total contraction time, total session time, total active time, rest time per repetition) were compared between conditions using paired-samples *t*-tests. Maximal torque was analyzed using a two-way repeated-measures analysis of variance (ANOVA) with Condition (with biofeedback vs. without biofeedback) and Time (pre-task vs. post-task) as within-subject factors. For all HDsEMG-derived variables (RMS, MNF, x-axis centroid, y-axis centroid, and modified entropy), two-way repeated-measures ANOVAs were conducted with Condition (with vs. without biofeedback) and Epoch (E1–E5) as within-subject factors. Borg CR10 ratings were compared between conditions using paired-samples t-tests (i) at task failure and (ii) at a matched workload, defined as the last repetition at which a rating was obtained in the shorter (no-biofeedback) session, compared with the rating given at the same repetition number in the biofeedback session. For the repeated-measures ANOVA analyses, when the assumption of sphericity was violated according to Mauchly’s test, Greenhouse–Geisser corrected p-values were applied. Post hoc pairwise comparisons for significant main effects or interactions were adjusted using Bonferroni correction. To characterize the time course of the medio-lateral centroid, its rate of change across epochs was additionally quantified for each participant and condition as the slope of a linear regression of the x-axis of the centroid against epoch number, and slopes were compared between conditions using a paired-samples t-test.

Within-participant changes across the task were quantified as the difference between Epoch 5 and Epoch 1 (ΔE5–E1) for each variable and condition. Condition differences were then calculated as the between-condition difference in these within-task changes (ΔE5–E1_biofeedback_ − ΔE5–E1_control_). Pearson correlation coefficients were computed between condition-related changes in the HDsEMG and torque variables (RMS, MNF, X-axis centroid, modified entropy, mean torque and torque CoV) and the endurance gain (Δrepetitions). Because the X-axis centroid and modified entropy showed a between-condition offset rather than a within-task change, their condition-related differences were additionally expressed as the mean over the five epochs (biofeedback − control). Additional correlations were performed between demographic variables (age, BMI, baseline MVC) and the individual response to biofeedback. Given the exploratory nature of these analyses and the small sample, correlations are reported with uncorrected p-values in the main text; Holm-adjusted p-values, 95% confidence intervals and the full correlation matrix are provided in the Supplementary Material (Tables S1–S2, Figure S1).

Effect sizes are reported as partial eta-squared (η*²_p_*) for ANOVA effects and Cohen’s *d* for paired comparisons. Statistical significance was set at α = 0.05 for all analyses.

## 3. RESULTS

### 3.1. Participant characteristics

Fourteen healthy adults (8 males, 6 females) completed both experimental sessions. Table 1 summarizes the participants’ anthropometric data (age: 31.0 ± 4.8 years; height: 171.6 ± 10.6 cm; body mass: 73.4 ± 16.8 kg; BMI: 24.7 ± 3.4 kg/m²) alongside key performance outcomes. Self-reported physical activity levels assessed via the IPAQ-LF (median 3711 [IQR 1458–4969] MET-min/week; Table 1) indicated that most participants were classified as highly active (n = 9, 64.3%), with four classified as moderately active (28.6%) and one as low active (7.1%).

**Table 1.** Participant characteristics, performance outcomes, self-reported physical activity level and perceived exertion. Values are mean ± SD, except IPAQ-LF, which is presented as median (IQR). Between-condition comparisons by paired t-tests. MVC was analyzed using a 2 × 2 (Condition × Time) repeated-measures ANOVA; the p value shown is for the Condition × Time interaction. *p < 0.05. BMI, body mass index; IPAQ-LF, International Physical Activity Questionnaire – Long Form; MET, metabolic equivalent of task; MVC, maximal voluntary contraction; CR10, Borg category-ratio 10 scale; IQR, interquartile range; SD, standard deviation; ANOVA, analysis of variance.

| ***Participant characteristics*** |  |  |  |
|---|---|---|---|
| n (male/female) | 14 (8/6) |  |  |
| Age (years) | 31.0 ± 4.8 |  |  |
| Height (cm) | 171.6 ± 10.6 |  |  |
| Body mass (kg) | 73.4 ± 16.8 |  |  |
| BMI (kg/m<sup>2</sup>) | 24.7 ± 3.4 |  |  |
| IPAQ-LF (MET-min/week) | 3711 (1458–4969) |  |  |
|  | **With Biofeedback** | **Without Biofeedback** | **p** |
| ***Performance outcomes*** |  |  |  |
| Number of repetitions | 19.2 ± 5.9 | 15.1 ± 4.1 | **<0.001\*** |
| Total contraction time (s) | 115.3 ± 35.6 | 90.4 ± 24.7 | **<0.001\*** |
| Total session time (s) | 119.6 ± 34.7 | 96.3 ± 26.9 | **<0.001\*** |
| Total active time (s) | 78.3 ± 22.9 | 64.3 ± 16.2 | **<0.001\*** |
| Rest time per repetition (s) | 2.18 ± 0.46 | 2.09 ± 0.46 | 0.381 |
| ***Maximal knee extension torque*** |  |  |  |
| Pre (N·m) | 253.9 ± 61.3 | 254.4 ± 58.1 | 0.258 |
| Post (N·m) | 187.6 ± 66.5 | 197.3 ± 59.7 |  |
| ***Perceived exertion*** |  |  |  |
| Borg CR10 at task failure | 9.00 ± 0.78 | 8.86 ± 1.03 | 0.336 |
| Borg CR10 at matched workload | 7.79 ± 1.48 | 8.86 ± 1.03 | 0.006* |

### 3.2. Maximal knee extension torque

The endurance task elicited significant neuromuscular fatigue in both sessions. The ANOVA revealed a significant main effect of Time for maximal torque (*F = 30.52, p < 0.001,* η*²_p_ = 0.701*; Figure 2), reflecting a substantial decline from pre-to post-task in both conditions. Importantly, the decline did not differ between conditions. There was no significant main effect of Condition (*F = 1.40, p = 0.258,* η*²_p_ = 0.097*), nor was there a Condition × Time interaction (*F = 1.40, p = 0.258,* η*²_p_ = 0.097*), indicating that MVCs were similar before and after the fatiguing task across conditions.

**Figure 2.**
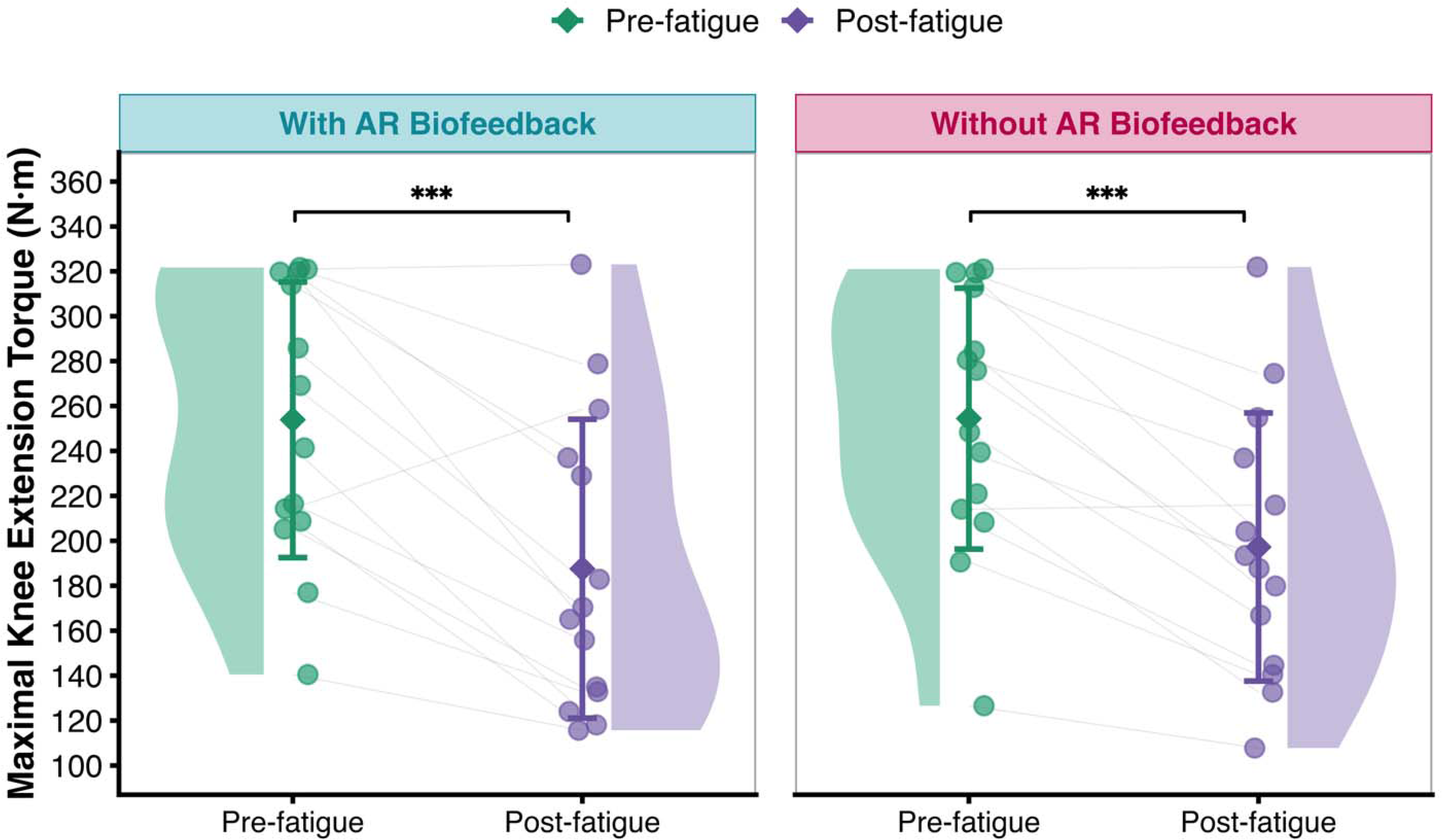
Maximal knee extension torque before and after the endurance task. Data distributions for maximum torque are visualized as raincloud plots (half-violin with individual data points) at pre- and post-fatigue timepoints. Green and purple diamonds denote the group mean (± SD) for the pre-task and post-task measurements, respectively, with paired lines connecting individual participant responses. Both the biofeedback and control conditions exhibited significant reductions in maximum torque following the endurance task (***p < 0.001); however, post-task maximum torque did not significantly differ between the two conditions (Condition × Time interaction, p = 0.258).

### 3.3. Rating of perceived exertion

The rating of perceived exertion (Borg CR10) at task failure was similar between conditions (p = 0.336; Table 1). However, when perceived exertion was compared at a matched workload (the final rated repetition of the no-biofeedback session), ratings were lower with AR biofeedback (7.79 ± 1.48 vs. 8.86 ± 1.03; t = −3.32, p = 0.006, d = 0.89; Table 1).

### 3.4. Torque output during the task

Despite the additional visual focus needed for the biofeedback condition, participants were able to adhere to the requested torque target, resulting in comparable torque characteristics across both trials. For mean torque, a significant main effect of Epoch was observed (*F = 6.19, p = 0.005,* η*²_p_ = 0.323*); Bonferroni-corrected pairwise comparisons showed that mean torque at Epoch 3 was significantly higher than at Epoch 4 (*p = 0.025*) and Epoch 5 (*p = 0.006*). However, no main effect of Condition (*F = 0.83, p = 0.380,* η*²_p_ = 0.060*) and Condition × Epoch interaction were observed (*F = 0.35, p = 0.739,* η*²_p_ = 0.027*) (Figure 3A).

**Figure 3.**
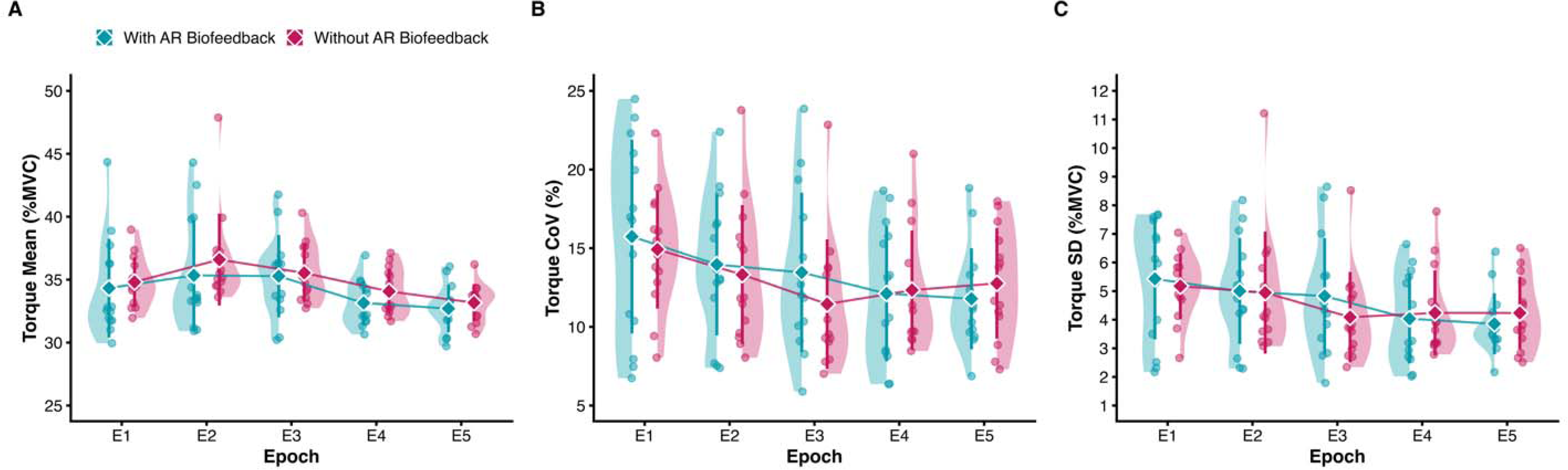
Mean torque and torque steadiness across epochs. Data distributions across five equal intervals (E1–E5) for (A) mean torque, (B) torque coefficient of variation (CoV), and (C) torque standard deviation (SD) are visualized as raincloud plots (half-violin with individual data points). Teal and pink diamonds denote the group mean (± SD) for the biofeedback and no-biofeedback conditions, respectively, connected by trend lines over time. All three metrics demonstrated a significant main effect of Epoch, with no significant differences observed between conditions.

Torque steadiness (CoV and SD of torque), likewise showed significant main effects of Epoch (CoV: *F = 4.37, p = 0.004,* η*²_p_ = 0.252*; SD: *F = 4.32, p = 0.004,* η*²_p_ = 0.250*); after correction, only torque CoV at Epoch 1 versus Epoch 4 differed significantly (p = 0.038). Neither of these metrics showed a significant main effect of Condition or Condition × Epoch interaction (*p > 0.05*) (Figure 3B, C).

### 3.5. Changes in endurance performance

Despite similar declines in maximum knee extension torque, comparable ratings of perceived exertion, and identical torque characteristics during the task, endurance performance was significantly greater in the condition of AR biofeedback. Participants completed 27% more repetitions with AR biofeedback than without biofeedback (*t = 5.29, p < 0.001, d = 1.41*; Table 1, Figure 4A). Similarly, total contraction time (*d = 1.41*), total session time (*d = 1.51*), and total active time (*d = 1.20*) were also significantly greater when AR biofeedback was provided (all *p < 0.001*). Importantly, this increase in work volume did not appear to reflect altered pacing, as the average rest time between repetitions did not differ between conditions (*p = 0.381*; Figure 4B).

**Figure 4.**
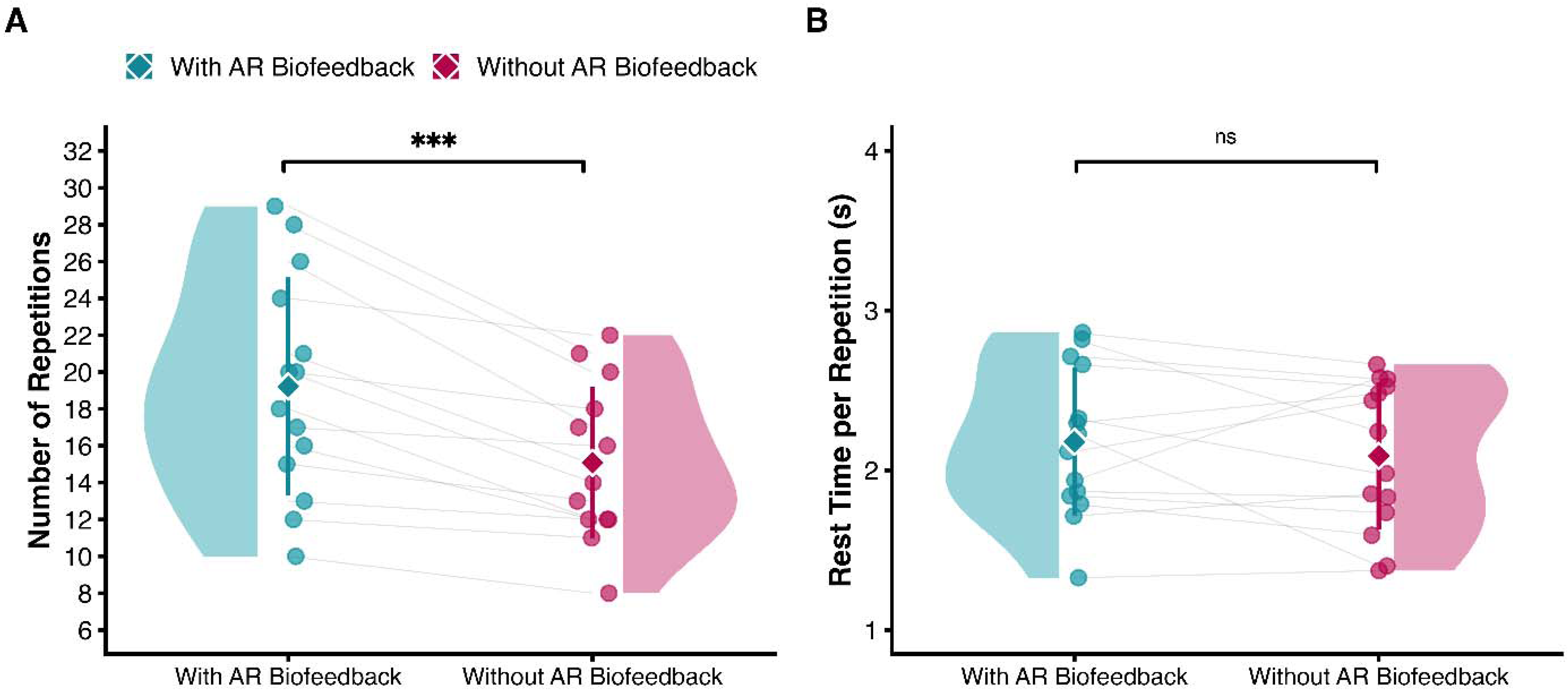
Effect of AR biofeedback on endurance performance. Data distributions for (A) the number of completed repetitions and (B) average rest time per repetition are visualized as raincloud plots (half-violin with individual data points). Teal and pink diamonds denote the group mean (± SD) for the biofeedback and no-biofeedback conditions, respectively, with paired lines connecting individual participant responses. A significant increase in repetitions was observed with biofeedback (***p < 0.001), whereas rest time per repetition did not differ between conditions (ns = not significant).

### 3.6. Changes in EMG variables

Indicators of myoelectric fatigue followed the expected trajectories throughout the task. RMS amplitude exhibited a progressive increase across epochs, with a significant main effect of Epoch (*F = 18.13, p < 0.001,* η*²_p_ = 0.582*; Figure 5A), whereas MNF decreased significantly over the same period (*F = 73.39, p < 0.001,* η*²_p_ = 0.850*; Figure 5B). The particularly large effect size observed for time on MNFs (η*²_p_ = 0.850*) confirms a robust spectral compression, consistent with progressive neuromuscular fatigue in both trials. Importantly, neither electromyographic metric was influenced by the addition of biofeedback. There were no significant main effects of Condition for either RMS (*F = 3.03, p = 0.105,* η*²_p_ = 0.189*) or MNF (*F = 0.003, p = 0.958,* η*²_p_ < 0.001*), nor were there any Condition × Epoch interactions (RMS: *F = 0.68, p = 0.539,* η*²_p_ = 0.050*; MNF: *F = 0.67, p = 0.532,* η*²_p_ = 0.049*).

**Figure 5.**
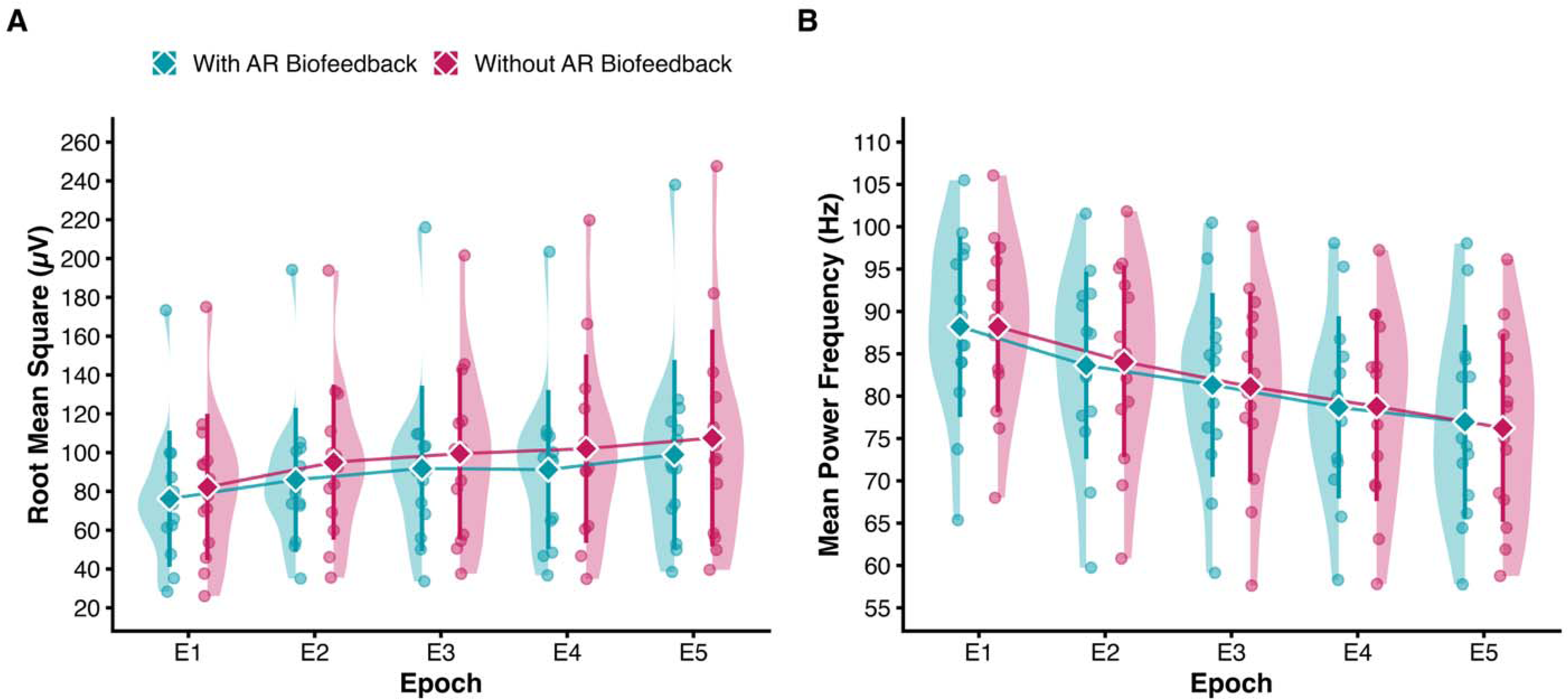
Myoelectric activity across epochs. Data distributions across five intervals (E1–E5) for (A) root mean square (RMS) amplitude and (B) mean power frequency (MNF) of knee extensor activity are visualized as raincloud plots (half-violin with individual data points). Teal and pink diamonds denote the group mean (± SD) for the AR biofeedback and no-biofeedback conditions, respectively, connected by trend lines over time. Both metrics demonstrated classic myoelectric manifestations of progressive fatigue, indicated by increasing RMS and decreasing MNF across epochs, with no significant differences observed between the two conditions.

### 3.7. Spatial distribution of muscle activity

The enhanced endurance performance observed during the AR biofeedback condition was accompanied by alterations in the spatial distribution of muscle activity. Specifically, the myoelectric activity was systematically located more laterally when AR biofeedback was provided. This was evidenced by a significant main effect of Condition for the medio-lateral axis (x-axis of the centroid) (*F = 6.81, p = 0.022,* η*²_p_ = 0.344*). No main effect of Epoch (F = 1.40, *p = 0.262,* η*²_p_ = 0.097*) or Condition × Epoch interaction (F = 0.75; *p = 0.501,* η*²_p_ = 0.055)*; Figure 6A) were observed. Per-participant linear slopes of the x-axis of the centroid across epochs did not differ between conditions (with: −0.19 ± 0.55 vs. without: −0.15 ± 0.49 mm/epoch; p = 0.683), indicating that the lateral offset was present from the first epoch and remained stable throughout the task.

**Figure 6.**
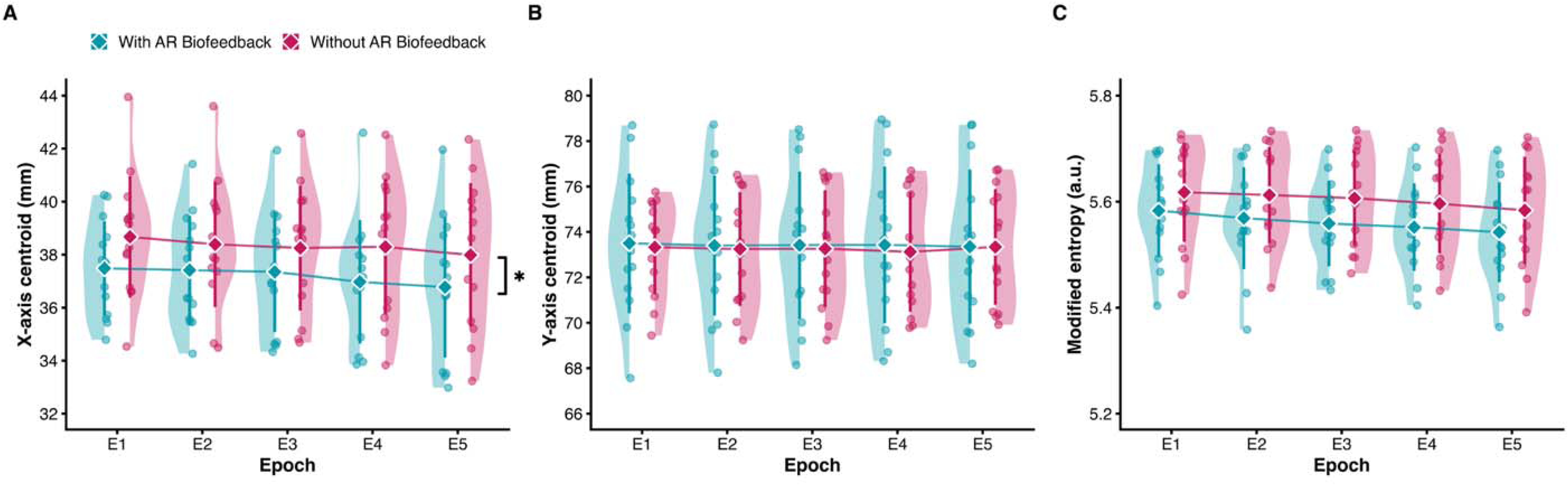
Spatial distribution of myoelectric activity across epochs. Data distributions across five intervals (E1–E5) for (A) the medio-lateral axis (X-centroid), (B) the proximo-distal axis (Y-centroid), and (C) modified entropy of knee extensor muscle activity are visualized as raincloud plots (half-violin with individual data points). Teal and pink diamonds denote the group mean (± SD) for the AR biofeedback and no-biofeedback conditions, respectively, connected by trend lines over time. The X-centroid exhibited a significant main effect of Condition (*p = 0.022), indicating that muscle activity was systematically located more laterally during the AR biofeedback trial. The Y-centroid remained unaffected, whereas modified entropy showed a non-significant tendency (p = 0.100) toward lower values, reflecting a more focalized and heterogeneous activation pattern with AR biofeedback.

In contrast, no changes were observed along the proximo-distal axis (Y-centroid) (Condition: F = 0.02, p = 0.891, η*²_p_ = 0.002*; Epoch: F = 0.18, p = 0.778, η*²_p_ = 0.014*), Condition × Epoch: F = 0.28, p = 0.767, η*²_p_ = 0.021*).

Finally, a non-significant tendency toward a main effect of Condition (*F = 3.14, p = 0.100,* η*²_p_ = 0.194*; Figure 6C) was observed for modified entropy. This trend was primarily driven by the biofeedback session, where lower entropy values reflected a more focalized and heterogeneous distribution of muscle activity. There were no observed main effects of Epoch (*F = 3.38, p = 0.077,* η*²_p_ = 0.206*) or Condition × Epoch interactions (*F = 0.27, p = 0.723,* η*²_p_ = 0.020*).

### 3.8. Exploratory correlations

Pearson correlations were utilized to assess relationships between condition differences in within-task EMG changes and the corresponding difference in the total number of repetitions performed. Condition-related changes in MNF (*r = −0.051, p = 0.863*) and RMS (*r = −0.002, p = 0.994*), as well as in centroid position, modified entropy, torque CoV and mean torque, showed no significant relationship with the performance improvement (all |r| ≤ 0.46, p ≥ 0.099). Furthermore, the AR biofeedback effect was not influenced by demographic or baseline characteristics such as age, BMI, or baseline MVC (all p > 0.05). Full results, including Holm-adjusted p-values and the same analysis with epochs defined over each condition’s own task duration, are given in the Supplementary Material (Tables S1–S2, Figure S1).

## 4. DISCUSSION

We investigated whether real-time AR HDsEMG biofeedback of knee extensor (VL and RF) HDsEMG could enhance endurance during dynamic fatiguing knee extension contractions. As hypothesized, participants receiving AR HDsEMG biofeedback completed significantly more repetitions before task failure. This improved performance was accompanied by an alteration in muscle activation patterns. Specifically, participants used the feedback to shift the center of their muscle activity laterally, as reflected by lower centroid values along the medio-lateral axis. Overall, these findings demonstrate that individuals can use real-time spatial feedback from HDsEMG to actively modify their motor strategy during a dynamic task, thereby improving their endurance.

### 4.1. Myoelectric manifestations of fatigue

As expected, global myoelectric indices of muscle fatigue (29) followed the well-established patterns during fatiguing contractions. RMS increased progressively, whereas MNF declined, consistent with previous studies (13, 30). Notably, both the magnitude and temporal progression of these responses were comparable across both conditions. When the two conditions were compared over the same absolute workload, the magnitude of these changes did not differ between conditions and was unrelated to the individual gain in repetitions, indicating that the additional work performed with AR biofeedback was not achieved by attenuating the development of myoelectric manifestations of fatigue (Supplementary Material, Tables S1–S2).

Similarly, no differences between conditions were observed in key physiological, perceptual, or behavioural markers of fatigue. The decline in maximal knee extension torque after the task, and the ratings of perceived exertion at task failure were comparable, indicating a similar level of physiological exhaustion. Torque steadiness and mean torque were likewise similar across conditions, suggesting that force control was not affected by the additional visual demands of the AR biofeedback. In addition, the average rest time per repetition did not differ, ruling out changes in pacing between conditions. Despite this overall similarity, participants using the AR HDsEMG biofeedback completed significantly more repetitions. This suggests that the improved endurance cannot be explained by differences in overall fatigue but is more likely due to local neuromuscular mechanisms that allowed greater mechanical work to be performed under the same conditions.

### 4.2. Mechanisms underlying the improvement of endurance

The spatial parameters offer important insight into potential mechanisms for improved performance. Throughout the AR biofeedback trial, the medio-lateral centroid was located more laterally, reflecting a greater engagement of the lateral portion of the knee extensors. Notably, this lateral offset was already present in the first epoch and remained stable across the task: the rate of change of the x-axis of the centroid across epochs was small and did not differ between conditions, indicating that participants adopted a more lateral activation pattern from the onset of the biofeedback trial rather than progressively shifting it as fatigue developed. As the largest quadriceps muscle (31), the VL possesses a large cross-sectional area and a substantial MU pool. By biasing activation toward this muscle, participants likely redistributed the load within the region covered by the grid, from the lateral portion of the RF toward the VL, and possibly also from muscles not recorded here (vastus medialis and the medial RF), thereby delaying task failure; the latter cannot be verified, as the grid covered only the VL and the lateral portion of the rectus femoris. This interpretation is consistent with the non-significant tendency toward lower entropy values during the AR HDsEMG biofeedback condition (*p = 0.100*), which indicates a more focal and region-specific activation pattern rather than a diffuse distribution of muscle activity across the electrode grid.

Nevertheless, the magnitude of this lateral shift did not directly correlate with the overall improvement in endurance. Additionally, the displacement was already clearly evident at the very onset of the task and did not increase as fatigue developed. This lack of direct correlation, coupled with the early adoption, suggests that the localized spatial reorganization was not a purely local adaptation to fatigue. Instead, this early shift might be better explained by other mechanisms, such as immediate biomechanical adjustments.

Visual inspection and post-task discussions supported this premise, revealing that participants frequently adopted a subtle internal hip rotation, often combined with a slight forward trunk lean, to view and alter the activation pattern displayed on their muscles; although this was not formally assessed or analyzed. This preference for internal over external rotation was likely dictated by the dynamometer’s positioning, which accommodated inward movement but restricted external rotation due to unwanted varus forces at the knee. We acknowledge that these postural adjustments were adopted only when the feedback was available and may therefore account for part of the between-condition difference; with the present design it is not possible to separate a change in the neural distribution of activity from a biomechanical adjustment prompted by the feedback (see Methodological considerations). This motor strategy offers a plausible biomechanical explanation for both the lateral centroid shift and the delayed task failure. Internal hip rotation alters the geometric orientation of the quadriceps compartments beneath the fixed electrode grid, mechanically biasing the amplitude distribution laterally. This is consistent with previous evidence that internal hip rotation is associated with increased VL activation (32). Additionally, the forward trunk lean increases hip flexion, shortening the biarticular rectus femoris and reducing its force-generating capacity through the length-tension relationship. To maintain the target torque, the nervous system may have to compensate by increasing drive to monoarticular synergists, particularly the VL. Although reducing the contribution of one synergist might be expected to hasten fatigue, transferring load to the muscle with the largest volume and MU pool of the group may increase the total force-generating capacity available for the task and thereby delay failure. Because the performance benefits may have also stemmed from these biomechanical adjustments rather than purely local neural adaptations within the muscle, this likely explains why the localized lateral shift alone did not correlate with the gains in endurance performance. Future studies incorporating kinematic recordings and multi-axis load cells to assess force direction are needed to test this hypothesis.

Beyond these individualized biomechanical adjustments, cognitive and attentional mechanisms likely contributed to the performance benefit. Participants reported equivalent perceived exertion at task failure and reached the same global fatigue, despite performing substantially more mechanical work with biofeedback. The AR display transformed internal neuromuscular information into a visual target in the participant’s environment, effectively functioning as an external attentional focus. Rather than consciously directing effort toward specific muscles, an internal focus that has been shown to constrain the motor system and reduce movement efficiency (33), participants monitored and responded to a real-time color map, which may have promoted more automatic motor control processes. This interpretation is consistent with evidence that an external focus of attention enhances muscular endurance during resistance exercise (34) and reduces muscle activity for equivalent force output (35). Additionally, the continuous, goal-directed nature of the feedback may have enhanced motivation and engagement, helping individuals sustain effort over a longer period. Consistent with this interpretation, when perceived exertion was compared at the same absolute workload (the last rated repetition of the control session), ratings were lower with AR biofeedback, whereas the myoelectric manifestations of fatigue over that same workload did not differ between conditions, suggesting that the additional repetitions were achieved through a perceptual and attentional benefit rather than through a slower development of peripheral fatigue. While the present design does not allow us to fully unravel the relative contributions of spatial reorganization, attentional redirection, and motivational engagement, all likely played a role in the observed endurance improvements. Regardless of the exact contribution of each mechanism, the successful use of this novel AR biofeedback to alter these parameters represents a significant advancement over how this technology is typically applied.

### 4.3. Technological and biomechanical mechanisms of spatial redistribution

To our knowledge, only two previous studies have attempted to alter the spatial activation of muscles using HDsEMG biofeedback. In our previous investigation (15), participants performed a sustained isometric shoulder abduction at 90° until task failure. While the visual feedback successfully induced a spatial shift in upper trapezius activity, it failed to improve endurance. This discrepancy is likely driven by both task constraints and local anatomy. The dynamic knee extension task in the current study provides greater biomechanical degrees of freedom, and the quadriceps’ substantially larger volume offers superior capacity for load redistribution. Conversely, Gaffney et al.(36) demonstrated that HDsEMG biofeedback successfully shifted trapezius activity toward the middle/lower regions during a typing task and this change was accompanied by adjustments in posture as shown by their kinematic data. This further supports our argument that biofeedback-guided spatial modifications may be intimately linked to broader biomechanical adjustments.

Beyond these mechanical factors, the method of feedback delivery likely played a vital role. Crucially, both prior investigations displayed the visual feedback on an abstract external monitor, requiring participants to cognitively translate spatially decoupled visual information onto their own movements. In contrast, our AR interface overlaid the biological information directly into the user’s physical environment, providing a highly intuitive and embodied experience that likely facilitated the rapid, successful adoption of their motor strategy. Together, these findings suggest that AR-projected HDsEMG biofeedback may have practical relevance beyond the laboratory, particularly in contexts where endurance performance, load distribution and motor relearning are central goals.

### 4.4. Methodological considerations

There are some methodological limitations that should be acknowledged. First, the relatively small sample size reduces statistical power for detecting interaction effects and may explain why some spatial metrics (e.g., modified entropy) only approached significance. Nevertheless, the performance effect was large and consistent across participants supporting its robustness. Second, HDsEMG-based spatial maps are inherently sensitive to subcutaneous tissue thickness, electrode-skin impedance, and potential crosstalk from adjacent muscles (37). Although the randomized crossover design mitigates systematic bias from these factors, and electrode placement was standardized using anatomical landmarks and skin markings, minor between-session spatial variability cannot be fully excluded. A further limitation is the absence of kinematic and multi-axis force measurements, which prevents precise quantification of postural or joint-level adjustments that may have contributed to the increased endurance. This is particularly relevant given evidence that voluntary modulation of MU activity within the quadriceps is constrained by common synaptic input and low-dimensional neural control structures (38). Consequently, the observed spatial redistribution may reflect a combination of neural and biomechanical factors, including posture, contraction mechanics, and force direction, rather than isolated voluntary control of muscle compartments (19, 38). Future studies integrating kinematic data and multi-axis force measurements are required to dissociate neural redistribution from compensatory biomechanical strategies. Additionally, the present analysis was limited to EMG amplitude distributions. MU decomposition was not performed, as the combination of dynamic contractions at 10°/s and progressive fatigue poses substantial challenges for current algorithms. Consequently, it remains unclear whether the observed spatial redistribution was driven primarily by neural adjustments, such as selective MU recruitment and rate modulation, or by biomechanical compensations. Future investigations using slower, non-fatiguing contractions that allow MU decomposition may better separate these specific mechanisms.

## 5. CONCLUSIONS

Novel AR HDsEMG biofeedback on the spatial distribution of knee extensor muscle activity can enhance endurance performance during dynamic fatiguing knee extension contractions, enabling individuals to modify the spatial pattern of muscle activation toward the lateral region of the knee extensors. In contrast to screen-based HDsEMG biofeedback, which altered muscle activation without improving endurance (15), projecting the map of the spatial distribution of muscle activity directly over the working muscle produced a 27% increase in repetitions under the same mechanical load. In athletic settings, this highlights its potential as a training tool to optimize neuromuscular load distribution in sports relying on repeated, high-force quadriceps contractions, such as cycling, skiing, sprinting and team sports. In rehabilitation, altered muscle activation patterns are common in musculoskeletal pain conditions, particularly in people with chronic low back pain (9, 12), and may contribute to rigid or maladaptive motor strategies; AR HDsEMG biofeedback may therefore provide a targeted method to retrain these patterns and restore more flexible neuromuscular control. Future studies are warranted to determine whether repeated exposure induces lasting motor adaptations in athletes and in patients with musculoskeletal pain.

## Supporting information

Suppementary Material

## Acknowledgments

The authors would like to thank all participants for their time and commitment to this study. This work was supported by the Royal Society (grant number IES\R3\223137).

## Data Availability

The datasets generated and analyzed during the current study are available from the corresponding author upon reasonable request.

## Author Contributions

M.A., G.L.C., M.G. and D.F. conceived and designed research; M.A. performed experiments; M.A. analyzed data; M.A., D.F., G.L.C. and M.G. interpreted results of experiments; M.A. prepared figures; M.A. drafted manuscript; M.A., G.L.C., M.G. and D.F. edited and revised manuscript; M.A., G.L.C., M.G. and D.F. approved final version of manuscript. D.F. is guarantor.

