## Supplementary material for "Real-time visualization of spatial muscle activity using augmented reality improves endurance during dynamic knee extensions": Suppementary Material

### **Supplementary File 1**

*Exploratory correlation analysis between the individual endurance gain and condition-related changes in HDsEMG, torque and participant characteristics (matched-workload epochs)*

#### **S1. Analysis overview**

For each participant, both experimental sessions were truncated to the number of repetitions completed in the shorter session (in every case the session without AR biofeedback) and the retained repetitions were divided into five equal epochs (E1–E5). All epoch-based variables below were therefore computed over the same absolute workload in the two conditions and refer to the combined electrode grid. Within-task changes were quantified as the difference between the last and first epoch (E5 − E1) in each condition; condition-related changes were then expressed as the between-condition difference of these within-task changes, ΔE5–E1_biofeedback_ − ΔE5–E1_control_, abbreviated ΔΔ. Because the medio-lateral (X-axis) centroid and modified entropy showed a tonic between-condition offset rather than a within-task change, they were additionally expressed as the mean over the five epochs, with − without. Pearson correlation coefficients were computed between each variable and the endurance gain (ΔReps). All correlations are exploratory (n = 14); p-values are reported uncorrected and, for completeness, after Holm adjustment across the 11 predictors tested against ΔReps (Table S1). Performance measures that are arithmetically related to ΔReps (total and active time, rest per repetition, repetitions per condition) are shown in the correlation matrix (Figure S1) for completeness but were not tested as predictors.

#### **S2. Variable definitions**

| **Abbreviation** | **Definition** | **Units** | **Sign** |
| --- | --- | --- | --- |
| Reps with / Reps without | Repetitions completed to task failure in each condition | n | — |
| ΔReps | Reps with − reps without biofeedback | n | + = more reps with biofeedback |
| ΔTotal time | Total session time, with − without | s | + = longer task with biofeedback |
| ΔActive time | Total active (contraction) time, with − without | s | + = longer with biofeedback |
| ΔRest/rep | Rest time per repetition, with − without | s | + = longer rests with biofeedback |
| ΔΔMNF | (E5 − E1) MNF, with − without | Hz | − = larger MNF decline with biofeedback |
| ΔΔRMS | (E5 − E1) RMS, with − without | µV | + = larger RMS increase with biofeedback |
| ΔΔTorque | (E5 − E1) mean torque, with − without | %MVC | − = larger torque decline with biofeedback |
| ΔΔCoV | (E5 − E1) torque CoV, with − without | % | + = less steady with biofeedback |
| ΔΔEntropy | (E5 − E1) modified entropy, with − without | a.u. | − = more focal with biofeedback |
| ΔΔX-centroid | (E5 − E1) X-centroid, with − without | mm | − = larger lateral drift with biofeedback |
| ΔX-centroid (mean) | Mean X-centroid over E1–E5, with − without | mm | − = more lateral with biofeedback |
| ΔEntropy (mean) | Mean modified entropy over E1–E5, with − without | a.u. | − = more focal with biofeedback |
| Baseline MVC | Highest pre-task MVC (session 1) | N·m | Participant characteristic |
| Age | Age at testing | years | Participant characteristic |
| BMI | Body mass index | kg/m² | Participant characteristic |

E1–E5, epochs 1–5 of the matched-workload window (both conditions truncated to the repetitions of the shorter session); (E5 − E1), within-task change; with − without, between-condition difference. MNF, mean power frequency; RMS, root mean square amplitude; CoV, coefficient of variation; MVC, maximal voluntary contraction; a.u., arbitrary units. Epoch-based variables were computed from the combined 10 × 6 differential grid.

#### **S3. Table S1. Correlates of the individual endurance gain (ΔReps)**

Pearson r with 95% confidence intervals (Fisher z transformation), uncorrected p and Holm-adjusted p (11 tests). Matched-workload (min-reps) epochs, n = 14.

| **Variable** | **r** | **95% CI** | **p (uncorrected)** | **p (Holm)** |
| --- | --- | --- | --- | --- |
| ΔΔMNF | −0.051 | −0.566 to 0.493 | 0.863 | 1.000 |
| ΔΔRMS | −0.002 | −0.532 to 0.529 | 0.994 | 1.000 |
| ΔΔTorque | −0.071 | −0.580 to 0.478 | 0.810 | 1.000 |
| ΔΔCoV | 0.017 | −0.518 to 0.543 | 0.953 | 1.000 |
| ΔΔEntropy | −0.458 | −0.795 to 0.096 | 0.099 | 1.000 |
| ΔΔX-centroid | 0.006 | −0.526 to 0.535 | 0.983 | 1.000 |
| ΔX-centroid (mean) | 0.177 | −0.390 to 0.647 | 0.545 | 1.000 |
| ΔEntropy (mean) | −0.124 | −0.614 to 0.435 | 0.673 | 1.000 |
| Baseline MVC | 0.165 | −0.400 to 0.640 | 0.572 | 1.000 |
| Age | −0.132 | −0.619 to 0.428 | 0.652 | 1.000 |
| BMI | −0.308 | −0.721 to 0.266 | 0.284 | 1.000 |

No variable was significantly associated with ΔReps. The largest coefficient was for ΔΔEntropy (r = −0.46, p = 0.099 uncorrected), which did not reach significance before or after adjustment.

#### **S4. Table S2. The same analysis using each condition's own endurance to define epochs (original analysis, for comparison)**

Epochs defined over each condition's full task (E5 = exhaustion in both conditions), as in the originally submitted manuscript. Shown to document that the association between ΔΔMNF and ΔReps reported previously (r = −0.592, p = 0.026, uncorrected) does not survive Holm adjustment and is absent when the two conditions are compared over the same absolute workload (Table S1), indicating that it reflected the longer task duration in the biofeedback condition rather than a difference in the rate of myoelectric fatigue.

| **Variable** | **r** | **95% CI** | **p (uncorrected)** | **p (Holm)** |
| --- | --- | --- | --- | --- |
| ΔΔMNF | −0.592 | −0.854 to −0.089 | 0.026 | 0.284 |
| ΔΔRMS | 0.474 | −0.075 to 0.803 | 0.087 | 0.866 |
| ΔΔTorque | −0.341 | −0.738 to 0.232 | 0.233 | 1.000 |
| ΔΔCoV | 0.169 | −0.397 to 0.642 | 0.564 | 1.000 |
| ΔΔEntropy | −0.397 | −0.766 to 0.169 | 0.160 | 1.000 |
| ΔΔX-centroid | 0.024 | −0.513 to 0.548 | 0.935 | 1.000 |
| ΔX-centroid (mean) | 0.084 | −0.468 to 0.588 | 0.776 | 1.000 |
| ΔEntropy (mean) | −0.210 | −0.667 to 0.360 | 0.470 | 1.000 |
| Baseline MVC | 0.165 | −0.400 to 0.640 | 0.572 | 1.000 |
| Age | −0.132 | −0.619 to 0.428 | 0.652 | 1.000 |
| BMI | −0.308 | −0.721 to 0.266 | 0.284 | 1.000 |

#### **S5. Figure S1 – Correlation Matrix**

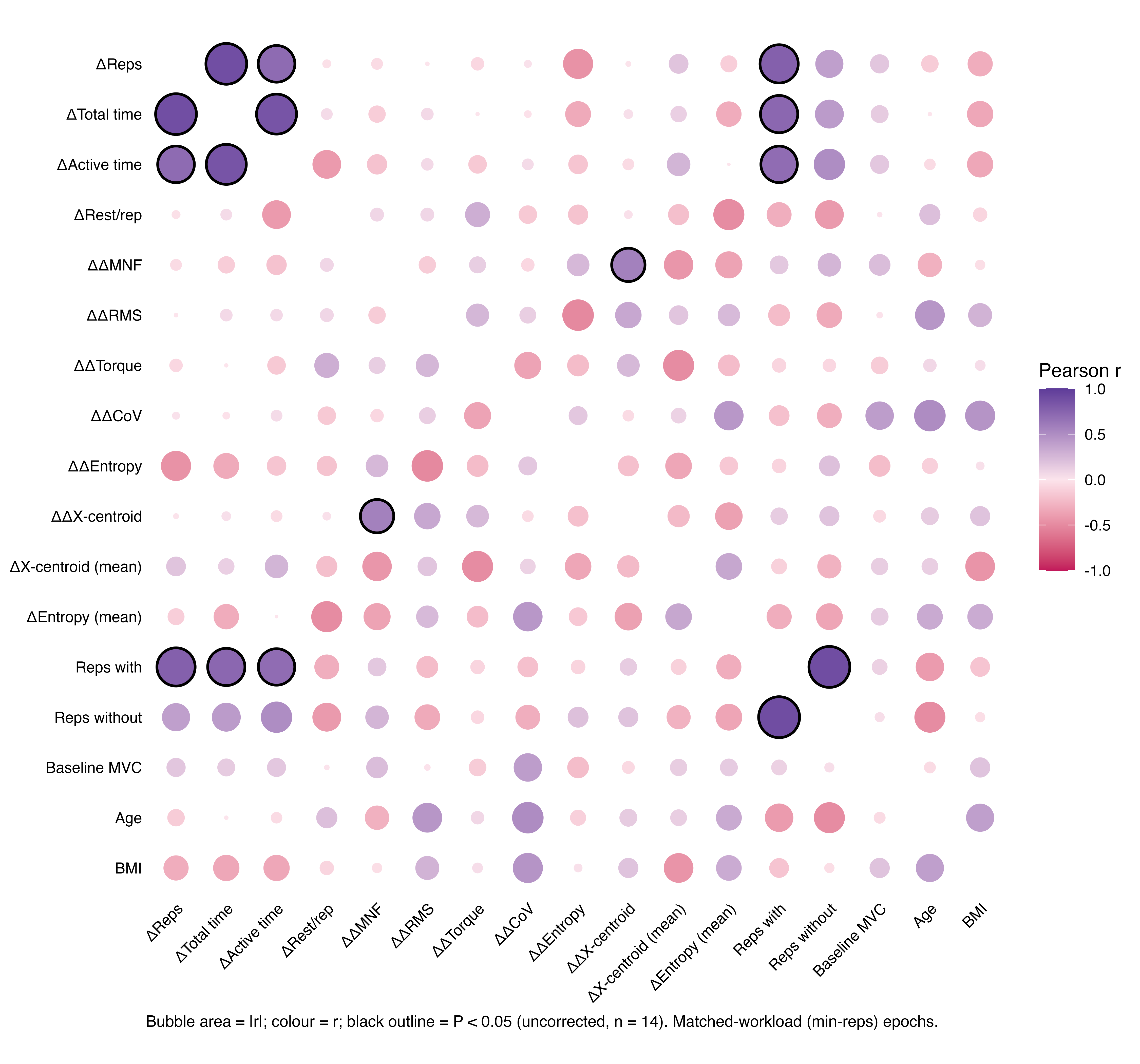

**Figure S1.** Correlation matrix (Pearson r) between the individual endurance gain (ΔReps), condition-related changes in the HDsEMG, torque and performance variables reported in the main text, and participant characteristics, computed over the matched-workload epochs (n = 14). Bubble area is proportional to |r|; colour encodes the sign and magnitude of r (magenta, negative; purple, positive); a black outline marks p < 0.05 (uncorrected). Variable definitions are given in Section S2. No HDsEMG, torque or participant variable was associated with ΔReps. The significant pairs involving ΔReps (ΔTotal time, ΔActive time, Reps with) are arithmetically related to it; the only significant pair among the HDsEMG predictors was ΔΔMNF with ΔΔX-centroid (r = 0.57, p = 0.033).
